# Supercharged PGR5-dependent cyclic electron transfer compensates for mis-regulated chloroplast ATP synthase

**DOI:** 10.1101/2022.09.25.509416

**Authors:** Gustaf E. Degen, Philip J. Jackson, Matthew S. Proctor, Nicholas Zoulias, Stuart A. Casson, Matthew P. Johnson

## Abstract

The light reactions of photosynthesis couple electron and proton transfers across the thylakoid membrane, generating NADPH, and proton motive force (pmf) that powers the endergonic synthesis of ATP by ATP synthase. ATP and NADPH are required for CO_2_ fixation into carbohydrates by the Calvin-Benson-Bassham cycle (CBBC). The dominant ΔpH component of the pmf also plays a photoprotective role in regulating photosystem II (PSII) light harvesting efficiency, through non-photochemical quenching (NPQ), and cytochrome *b*_6_*f* (cyt*b*_6_*f*) to photosystem I (PSI) electron transfer, via photosynthetic control. ΔpH can be adjusted by increasing the proton influx into the thylakoid lumen via upregulation of cyclic electron transfer (CET) or decreasing proton efflux via downregulation of ATP synthase conductivity (gH^+^). The interplay and relative contributions of these two elements of ΔpH control to photoprotection are not well understood. Here, we show that an Arabidopsis ATP synthase mutant (*hope2*) with 40% higher proton efflux, has supercharged CET. Double crosses of *hope2* with the CET-deficient *pgr5* and *ndho* lines reveal that PGR5-dependent CET is the major pathway contributing to higher proton influx. PGR5-dependent CET allows *hope2* to maintain wild-type levels of ΔpH, CO_2_ fixation and NPQ, however photosynthetic control remains absent, and PSI is acceptor-side limited. Therefore, high CET in the absence of ATP synthase regulation is insufficient for PSI photoprotection.

## Introduction

CO_2_ fixation into biomass during photosynthesis requires reducing power in the form of NADPH and energy in the form of ATP (Kramer and Evans, 2010). NADPH is provided by coupled photosynthetic linear electron transfer (LET) reactions in the thylakoid membrane, which also generate pmf for ATP synthesis via ATP synthase. In chloroplasts, pmf is largely composed of the proton concentration gradient (ΔpH), with minimal contribution from the membrane potential (ΔΨ) (Wilson et al., 2021), which is detrimental to productive charge separation in PSII (Davis et al., 2016) and largely dissipated by counterion movements (Hind et al., 1974). In addition to its manifest role in ATP synthesis, the ΔpH also plays a vital role in regulating photosynthetic electron transfer and light harvesting reactions via photosynthetic control and energy-dependent NPQ, known as qE (Li et al., 2009; Malone et al., 2021). Photosynthetic control restricts the rate of plastoquinol (PQH_2_) oxidation at cyt*b*_6_*f* activity, can be measured as the donor-side limitation of PSI (Y(ND)) using P700 absorption spectroscopy and protects PSI against photo-oxidative damage in excess light (Jahns et al., 2002; Suorsa et al., 2013). In contrast, qE involves ΔpH induced protonation of the PsbS protein and de-epoxidation of the light harvesting antenna complex II (LHCII)-bound xanthophyll violaxanthin to zeaxanthin, which collectively bring about energy dissipation in LHCII, protecting PSII from photo-oxidative damage (Ruban et al., 2012). qE can be measured as the rapidly-relaxing component of NPQ using pulse-amplitude modulated chlorophyll fluorescence. These ΔpH-dependent regulatory mechanisms are critical to plant growth in fluctuating light environments and rely on the careful modulation of the proton influx/ efflux reactions across the thylakoid membrane (Armbruster et al., 2017).

Proton efflux is regulated primarily by the conductivity (gH^+^) and abundance of the chloroplast ATP synthase (Kramer et al., 2004). Antisense mutants of the γ-subunit in tobacco showed that ATP synthase levels could be reduced by 50% without affecting gH^+^ or pmf, while further decreases caused decreased gH^+^ and increased pmf leading to higher qE and Y(ND) (Rott et al., 2011). Regulation of the ATP synthase activity is therefore a key element of proton efflux control. Two types of regulation have been described for ATP synthase; redox and metabolic control (Mills and Mitchell, 1982; Ort and Oxborough, 1992; Kanazawa and Kramer, 2002; Kohzuma et al., 2013). Redox control of ATP synthase is mediated by the reduction-oxidation status of two regulatory cysteines (C202, C208 in Arabidopsis) which form a disulfide bridge stabilising a loop of the γ1-subunit that acts as a chock, interfering with the rotation of the catalytic F1 head of the enzyme involved in ATP synthesis (Hisabori et al., 2003; Hahn et al., 2018). Therefore a lower threshold pmf is required to activate the reduced enzyme (Junesch and Gräber, 1987). Upon illumination, activation of LET causes reduction of Fd and NADPH, these can reduce thioredoxin (TRX) proteins via the Fd dependent thioredoxin reductase (FTR) or NADPH dependent thioredoxin reductase (NTRC) enzymes. TRX then reduces the regulatory disulfide in the γ1-subunit (Carrillo et al., 2016; Sekiguchi et al., 2020). In contrast, inactivation of LET in the dark leads to gradual oxidation of the regulatory cysteines by 2-Cys peroxiredoxin (PRX), restoring the higher pmf threshold for activation (Ojeda et al., 2018). In addition to redox control, gH^+^ is known to be modified by varying CO_2_ and Pi concentration suggesting ATP synthase also responds to the metabolic state of the stroma (Kanazawa and Kramer, 2002; Avenson et al., 2005; Takizawa et al., 2008; Kohzuma et al., 2013). The *mothra* mutant of Arabidopsis has changes in three conserved acidic residues in the γ1-subunit (D211V, E212L and E226L) resulting in the loss of redox sensitivity, yet metabolic control is unaffected. The pmf threshold for activation in *mothra* is correspondingly higher resulting in a lower gH^+^, increased pmf, increased qE and lower LET rate (Kohzuma et al., 2012). In contrast, in the Arabidopsis γ1-subunit mutant *hope2* (<u>h</u>unger for <u>o</u>xygen in <u>p</u>hotosynthetic <u>e</u>lectron transport <u>2</u>, G134D), ATP synthase gH^+^ was insensitive to changing CO_2_ concentration indicating metabolic control was lost, although redox control was reportedly unaffected (Takagi et al., 2017). *Hope2* showed the opposite phenotype to *mothra* with increased gH^+^, the virtual absence of Y(ND) and a greater susceptibility to PSI photoinhibition, although maximum LET rate and CO_2_ assimilation were unaffected. Crucially, the phenotype of *hope2* was successfully ameliorated via complementation with a WT copy of the γ1-subunit.

Proton influx can occur via one of several coupled electron transfer pathways. LET involves the light-powered transfer of electrons from water to NADP^+^, via a chain including PSII, plastoquinone (PQ)/PQH_2_, cyt*b*_6_*f*, plastocyanin (Pc), PSI, ferredoxin (Fd) and ferredoxin-NADP^+^ reductase (FNR). Unlike LET, alternative electron flows can contribute to pmf generation without generating net NADPH. These include; pseudo-cyclic electron transfer (water-water cycle), where electrons from Fd are instead transferred to oxygen via flavodirron (Flv) proteins to form water; the Mehler reaction, where PSI directly reduces oxygen to superoxide; and the malate valve, where NADPH is consumed to reduce oxaloacetate to malate, which can be exported from the chloroplast to be oxidised in the mitochondria (Miyake, 2010; Alric and Johnson, 2017). However in angiosperms such as Arabidopsis, Flv proteins are absent and the primary alternative electron flow is cyclic electron transfer (CET), where electrons from Fd reduce PQ forming a cycle around PSI and cyt*b*_6_*f* via Fd-PQ reductase activity (FQR) (Johnson, 2011; Yamori and Shikanai, 2015). Two CET pathways occur in Arabidopsis, one sensitive to the inhibitor antimycin-A (AA) involves the Proton Gradient Regulation 5 (PGR5) protein (referred to as CET1) and the second is catalysed by the NDH-like photosynthetic Complex I (NDH, referred to as CET2) (Yamori and Shikanai, 2015). How PGR5 mediates CET1 remains unknown, early ideas that it acts together with PGRL1 to form an FQR (DalCorso et al., 2008; Hertle et al., 2013) were recently invalidated by new evidence showing that PGRL1 is not essential (Rühle et al., 2021). Instead, PGRL1 binds PGR5 to protect it from degradation by PGRL2, and therefore CET1 can function if both PGRL1 and PGRL2 are absent. An alternative suggestion is that the cyt*b*_6_*f* complex binds FNR and together they play the role of the FQR (Shahak et al., 1981; Joliot and Johnson, 2011). However while cyt*b*_6_*f* can be co-purified with FNR (Clark et al., 1984; Zhang et al., 2001), to date no FNR-cyt*b*_6_*f* -PGR5 complex possessing the requisite FQR activity, which can be as high as 130 e^-1^ s^-1^ in Arabidopsis (Joliot et al., 2004), has been isolated. In contrast, high -resolution structures of the NDH-PSI CET2 supercomplex from Arabidopsis and barley have been described (Shen et al., 2022; Su et al., 2022). The Arabidopsis *ndho* and *crr* mutants, which both lack NDH-dependent CET2, have relatively mild phenotypes with only slight differences seen in pmf generation and photosynthetic activity (Munekage et al., 2004; Wang et al., 2015; Nikkanen et al., 2018). On the other hand, the Arabidopsis *pgr5* mutant suffers a significant loss of ΔpH, qE and Y(ND) in high light together with lower LET, CO_2_ assimilation and increased PSI photoinhibition (Munekage et al., 2004; Suorsa et al., 2012; Nikkanen et al., 2018). The more severe phenotype of *pgr5* suggests that CET1 is the dominant pathway in Arabidopsis and that NDH-dependent CET2 has a limited capacity to compensate. A number of high cyclic electron flow (*hcef)* mutants have been described in Arabidopsis and tobacco (Livingston et al., 2010a; Livingston et al., 2010b; Strand et al., 2017). Yet to date, those *hcef* mutants characterised in detail have only shown upregulation of the NDH-dependent CET2 pathway (Livingston et al., 2010a; Livingston et al., 2010b; Strand et al., 2017), leading some to speculate that PGR5 may not be directly involved in CET (Suorsa et al., 2012; Takagi and Miyake, 2018). Interestingly, *pgr5* also shows a high gH^+^ phenotype, leading to the suggestion that it may alternatively regulate ATP synthase (Avenson et al., 2005).

The fact that *hope2* and *pgr5* mutants share a low Y(ND), high gH^+^ phenotype but differ in their respective capacities for qE, CO_2_ assimilation and LET (Munekage et al., 2004; Takagi et al., 2017) suggests that proton influx and efflux may play distinct roles in photosynthetic regulation. Here, we investigated these relationships further by creating double mutants lacking *hope2* and either *ndho* or *pgr5*. The results unexpectedly demonstrate that loss of ATP synthase regulation in *hope2* is compensated for by increased PGR5-dependent CET, which maintains ΔpH and qE but fails to restore photosynthetic control.

## Results

### *hope2* maintains wild-type levels of pmf due to increased proton flux

We first sought to confirm that the CO_2_ assimilation (A) phenotype of *hope2* was WT-like as previously reported (Takagi et al., 2017). Indeed, A in *hope2* was similar to WT Col-0 at both high and low light intensity, although between 100 and 275 µmol photons m^-2^ s^-1^ A was slightly lower in *hope2* (Figure 1a). The A/Ci response in *hope2* was not significantly different to the WT Col-0 (Figure 1b). In contrast, in *pgr5* we find A in response to light and varying intercellular CO_2_ concentrations (Ci) is lower compared to the WT *gl1* (Figure 1a and b). To estimate maximum Rubisco carboxylation rates *in vivo* (V_c,max_) and maximum electron transport rate used in RuBP regeneration (J_max_), we fit the Farquhar-von Caemmerer-Berry (FvCB) model (Farquhar et al., 1980) to individual A/Ci curves. V_c,max_ and J_max_ were not different in *hope2* compared to Col-0 (p > 0.05), confirming that carbon fixation is not limited, in contrast to CET1-deficient *pgr5* (p < 0.05). This confirms that despite sharing the high gH^+^ phenotype with *pgr5, hope2* is still able to maintain an optimal ATP/NADPH ratio for CO_2_ assimilation.

**Figure 1:**
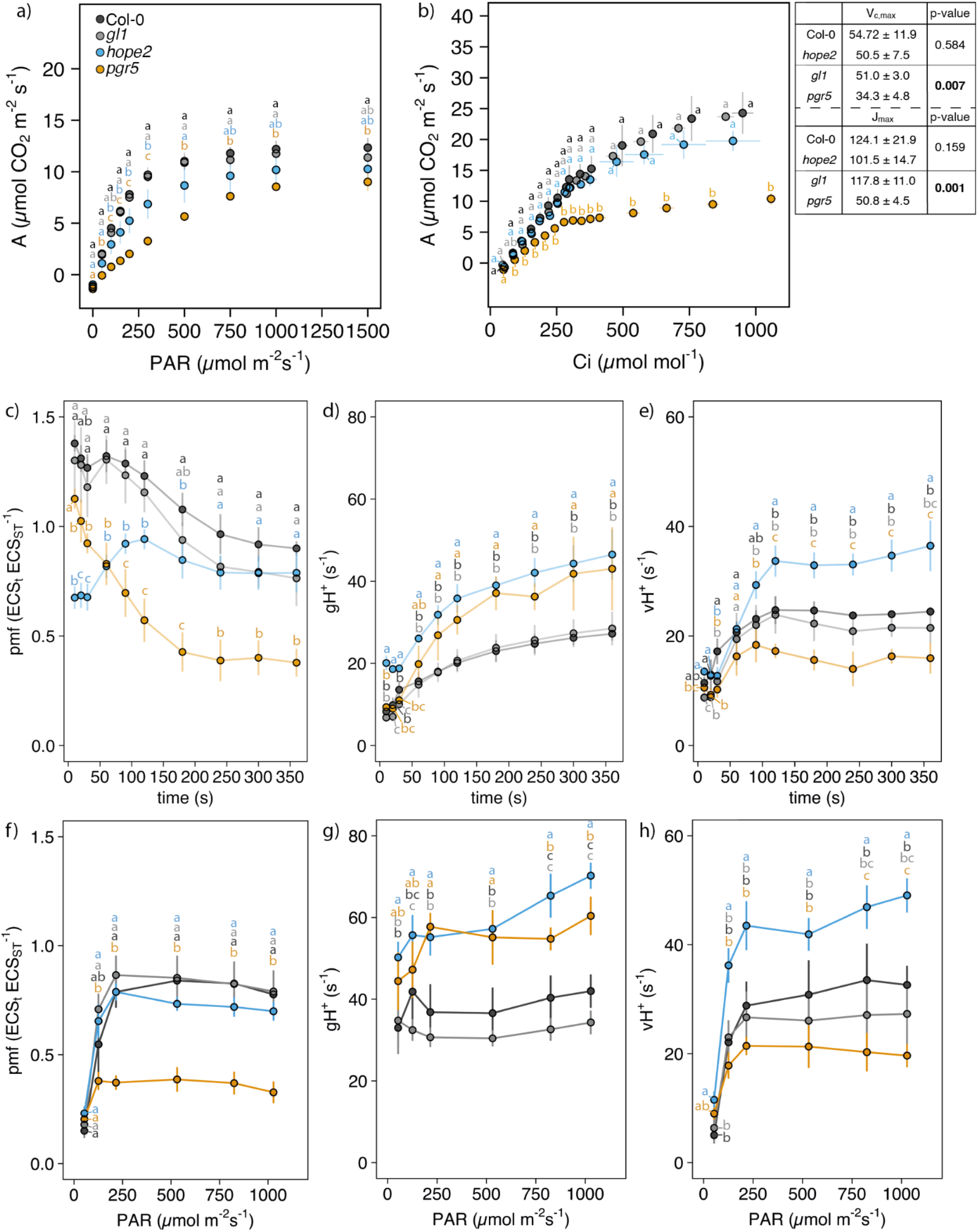
Gas-exchange and ECS measurements of WT, *hope2* and *pgr5*. a) CO_2_-assimilation at various light intensities. Plants were adapted to 400 ppm reference CO_2_ and 1500 µmol photons m^-2^ s^-1^ light for at least 10 min until steady-state was reached. Light levels were decreased stepwise, and data logged after at least 3 min at each light intensity. b) CO_2_-assimilation at various Ci concentrations. Plants were adapted to 400 ppm reference CO_2_ and 1500 µmol photons m^-2^ s^-1^ light for at least 10 min until steady-state was reached. Reference CO_2_ was decreased to 50 ppm, increased to 400 ppm and finally increased to 1250 ppm. Data was logged after 90 – 120 s at each CO_2_ concentration. Table: Maximum Rubisco activity (V_c,max_) and maximum electron transport rate used in RuBP regeneration (J_max_). c, f) proton motive force (pmf); d, g) proton conductance of ATP synthase (gH^+^); e, h) proton flux (vH^+^). c-e) Measurements were taken during photosynthetic induction at the indicated time points using 169 µmol photons m^-s^ s^-1^ actinic light. f-h) Measurements were taken after 5 min at each indicated light intensity. Data points represent the average of 3-7 biological replicates ± SD. Colours represent genotypes analysed: Col-0 (black), *gl1* (grey), *hope2* (blue) and *pgr5* (orange). Different letters indicate statistical significance between genotypes at each time point, light intensity or Ci, calculated from a Tukey HSD test, alpha = 0.05. Bold p-values in (b) indicate significantly different V_c,max_ and J_max_ between Col-0 and *hope2*, calculated from a two-sided t-test, alpha = 0.05.

We investigated how *hope2* is able to achieve WT-like CO_2_ assimilation further by comparing the generation of pmf in *hope2* and *pgr5* during photosynthetic induction. During the first 50 s of illumination, pmf in *hope2* was lower compared to *pgr5* and WT (Figure 1c), due to increased gH^+^ levels (Figure 1d), while proton flux (vH^+^) was similar to *pgr5* and WT (Figure 1e). Thus, during the first 50 s of photosynthetic induction gH^+^ regulation makes a larger contribution to pmf than vH^+^. After 3 min, pmf in *hope2* reached WT levels, despite high gH^*+*^, due to strongly increased vH^+^ (Figure 1c-e). In contrast, pmf in *pgr5* dropped continuously during the first ∼200 s of actinic light exposure due to a combination of increasing gH^+^ and low vH^+^ (Figure 1c-e). Therefore, on longer timescales increases in vH^+^ are important for maintaining pmf in the WT as the gH^+^ regulation relaxes. In *hope2*, the lack of gH^+^ regulation leads to a compensatory increase in vH^+^, maintaining pmf at WT levels beyond ∼150 s illumination. Having established that *hope2* had WT-level pmf after 6 min of low actinic light, we next investigated how *hope2* and *pgr5* behaved during increasing light intensities. This revealed that pmf in the WT plateaued at ca. 0.8 at 250 µmol photons m^-2^ s^-1^ (Figure 1f), similar to previous reports (Nishikawa et al., 2012; Wang et al., 2015; Yamamoto et al., 2016; Nikkanen et al., 2018; Nikkanen et al., 2019; Yamamoto and Shikanai, 2020; Hepworth et al., 2021; Rühle et al., 2021) and pmf in *hope2* was not significantly different at all light intensities. gH^+^ and vH^+^ on the other hand, were still higher in *hope2* (Figure 1g and h), confirming that increased vH^+^ drives the maintenance of WT levels of pmf in *hope2*. On the other hand, *pgr5* had diminished pmf at all but the lowest light intensity due to a combination of lower vH^+^ and higher gH^+^ compared to the WT (Figure 1g and h).

### Cyclic electron transfer is upregulated in *hope2*

We find that increased gH^+^ in *hope2* is compensated for by an increase in vH^+^, which could be caused by either increased LET, CET or another alternative electron flow. To test this further, we compared the thylakoid proteomes for proteins that were up or downregulated in *hope2* relative to the WT (Supplemental Figure S1). As expected, ATP synthase abundance was decreased by ∼50% in *hope2* consistent with past immunoblotting results (Takagi et al., 2017). Other proteins significantly upregulated (p < 0.05) included the state transition kinases STN7 and STN8, PsbS, violaxanthin de-epoxidase (VDE), the H^+^/K^+^ antiporter KEA3, while those downregulated included TRXM2, PSAO and PSBT. Most strikingly however, PGR5 (0.56-fold increase) and multiple subunits of the NDH complex (ndhF, H, I, K, M, N, O, S and U) (∼0.21-0.41-fold increase) showed significantly increased abundance in *hope2* (Figure 2a). Interestingly however, PGRL1 was unchanged in *hope2* (Figure 2a). Increased abundance in *hope2* was also seen for LFNR1 and LFNR2 and their membrane tether, thylakoid rhodanese-like protein (TROL), which have been recently linked to CET (Kramer et al., 2021). On the other hand, in *pgr5*, both PGRL1 isoforms and NDH subunits were decreased by ∼0.70-1.31 and ∼0.21-0.35-fold respectively (Figure 2a), as shown previously by immunoblotting (Munekage et al., 2004; Nikkanen et al., 2018; Wada et al., 2021). Given these results we hypothesised that pmf may be maintained in *hope2* via increased CET relative to WT. A well-established method of assessing changes in CET is the relationship between vH^+^ and LET (calculated by chlorophyll fluorescence) (Okegawa et al., 2005; Livingston et al., 2010a; Livingston et al., 2010b; Strand et al., 2015; Strand et al., 2017). A steeper slope indicates a greater contribution of CET to vH^+^. In *hope2*, the relationship between vH^+^ and LET was significantly (p < 0.05) steeper than in the WT Col-0, with a slightly shallower slope observed in *pgr5* (Figure 2b). We estimate from the slope that vH^+^ is decreased by 7.9% in *pgr5*, similar to the 13% previously determined by this method (Avenson et al., 2005), but increased by 48% in *hope2*. Deviation of a linear relationship between the quantum yields of PSI (Y(I)) and PSII (Y(II)) at varying light intensities provides another indication of CET capacity (Okegawa et al., 2005; Livingston et al., 2010a; Livingston et al., 2010b; Strand et al., 2015; Strand et al., 2017). Consistent with higher CET in *hope2* we found increased Y(I) relative to Y(II) compared to the WT, while the smallest deviation was found in *pgr5* (Figure 2c). An alternative complementary method of assessing CET *in vivo* involves following the rate of P700 oxidation induced by far red (FR) light which preferentially excites PSI (Joliot and Joliot, 2002). In this assay, decreased CET activity results in faster P700 oxidation and lower half-times (t_0.5_), whereas increased CET has the opposite effect (Joliot and Joliot, 2002; Joliot and Johnson, 2011; Rühle et al., 2021). Consistent with lower CET in *pgr5*, FR-light induced P700 oxidation was faster than WT *gl1* (Figure 2d). However, P700 oxidation t_0.5_ in *hope2* was significantly (p = 0.0200) slower than Col-0 (Figure 2d), in line with increased CET relative to WT Col-0. We confirmed that these differences in P700 oxidation rate could not be ascribed to differences in antenna size between the WT Col-0 and *hope2* by infiltration of leaves with DCMU and methyl viologen, which eliminate donor and acceptor side limitations on PSI (Supplemental Figure S2).

**Figure 2:**
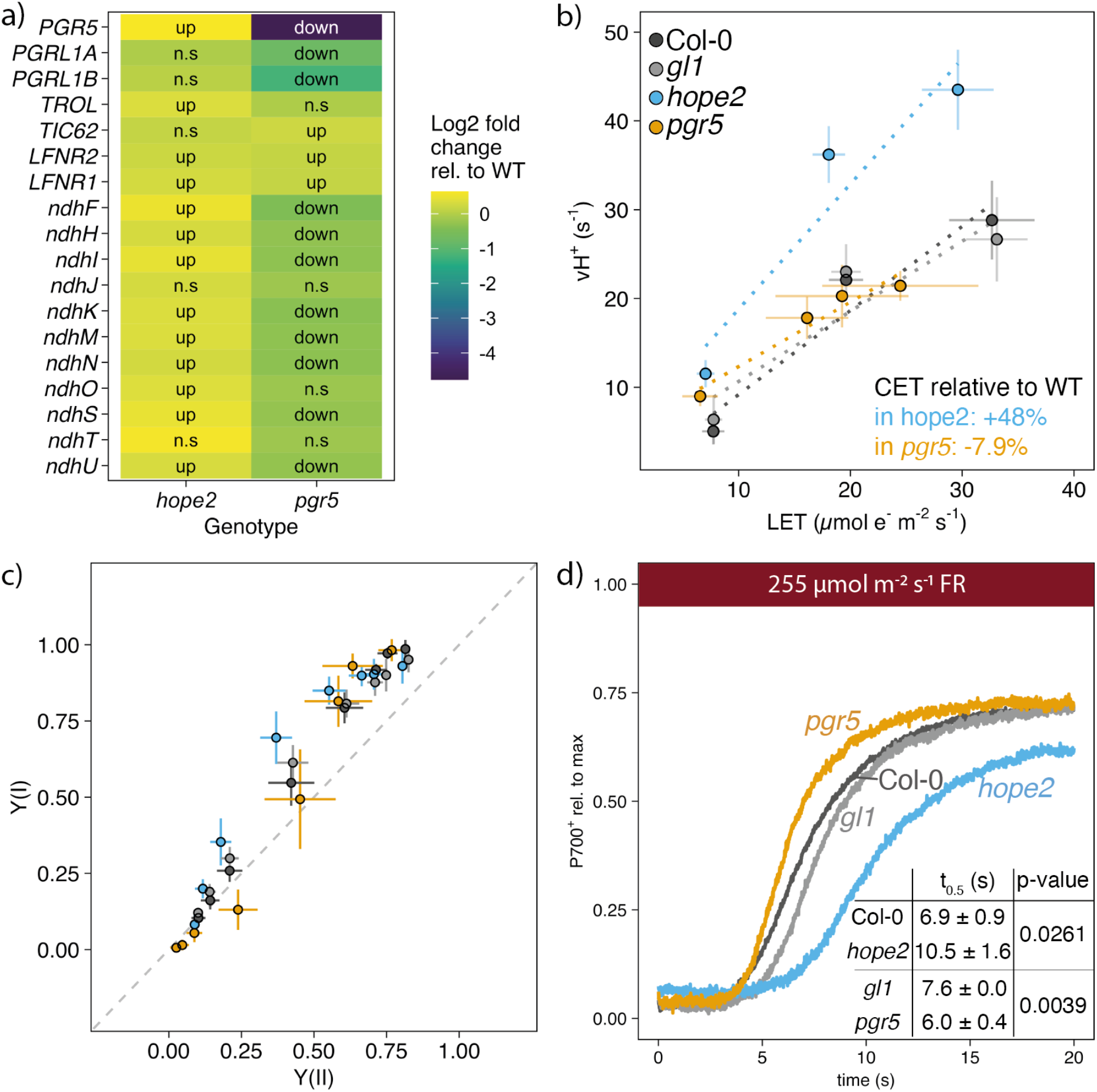
Measurements of cyclic electron transfer. a) Abundance of proteins involved in CET, normalised to WT. b) Proton flux vs. linear electron transfer. Higher flux at similar LET indicates increased cyclic electron transfer. c) Yield of photosystem I vs. yield of photosystem II. d) P700 oxidation during FR light. Prior to FR light, leaves were exposed to a weak ML for 30 s, followed by a SP and 30 s of darkness. Data was normalised to maximal P700 oxidation after 30 s FR light and a SP. Insert: Half-life of P700 oxidation determined by fitting an allosteric sigmoidal function prior to the SP. Data points represent the average of 3-6 biological replicates ± SD. Colours represent genotypes analysed: Col-0 (black), *gl1* (grey), *hope2* (blue) and *pgr5* (orange). Different letters indicate statistical significance between genotypes at each time point, calculated from a Tukey HSD test, alpha = 0.05. In (a) “Up” indicates significantly (p<0.05) more abundant proteins and “down” significantly (p>0.05) less abundant proteins.

### PGR5 is the major CET pathway in *hope2*

Since both NDH and PGR5 abundance increased in *hope2* we sought to ascertain whether one or both pathways contributed to the observed increases in CET and vH^+^. To that end the double mutants *hope2 ndho* and *hope2 pgr5* were created and verified by DNA sequencing for the *hope2* mutation and immunoblotting for NdhS and PGR5 levels (Supplemental Figure S3). Previous high CET mutants have involved the NDH pathway (Livingston et al., 2010a; Livingston et al., 2010b; Strand et al., 2015; Strand et al., 2017), which prompted us to first analyse the *hope2 ndho* double mutant. At light levels above 250 µmol photons m^-2^ s^-1^, pmf in the *hope2 ndho* double mutant was not significantly different to the *hope2* and *ndho* single mutants (Figure 3a), though both *ndho* and *hope2 ndho* showed slightly lower pmf than the WT, consistent with past results (Nikkanen et al., 2018). Similarly, gH^+^ and vH^+^ were largely unchanged in *hope2 ndho* compared to *hope2*, which were both significantly higher than *ndho* and WT (Figure 3b-c). The NPQ level in the *hope2* mutant was WT-like, whereas *ndho* showed higher NPQ as previously reported (Rumeau et al., 2005; Takagi et al., 2017). The *hope2 ndho* mutant showed an NPQ level between *hope2* and *ndho*, demonstrating that NPQ in *hope2* does not require NDH (Figure 3d), although clearly elevated NPQ in *ndho* is affected by loss of gH^+^ regulation. The PSII quantum yield (Y(II)) and PSII Q_A_^-^ reduction (1-qL) were similar in *hope2 ndho* and *hope2*, only low light had a moderate effect (Figure 3e-f). The PSI quantum yield (Y(I)) is increased in *hope2* relative to the WT at moderate light intensities between 250 and 500 µmol photons m^-2^ s^-1^, whereas *hope2 ndho* and *ndho* were lower than WT and *hope2* at low light but was similar at higher light intensities (Figure 3g). Finally, PSI donor side limitation (Y(ND)) and PSI acceptor-side limitation (Y(NA)) in *hope2 ndho* were also similar to *hope2*, which were significantly lower than the WT and *ndho* (Figure 3h-i). CET measured via the relationship between vH^+^ and LET or Y(I) and Y(II) in *ndho* was similar to WT, whereas *hope2 ndho* behaved largely like *hope2*, with only slight deviation seen at the lowest LET levels (Figure 3j,k), suggesting the major contribution of the NDH pathway is in low light. Furthermore, CET measured via the P700 oxidation method in *hope2 ndho* showed it was unchanged compared to *hope2* (p = 0.2424), suggesting the elevated CET is little affected by the absence of the NDH pathway under FR illumination (Figure 3l).

**Figure 3:**
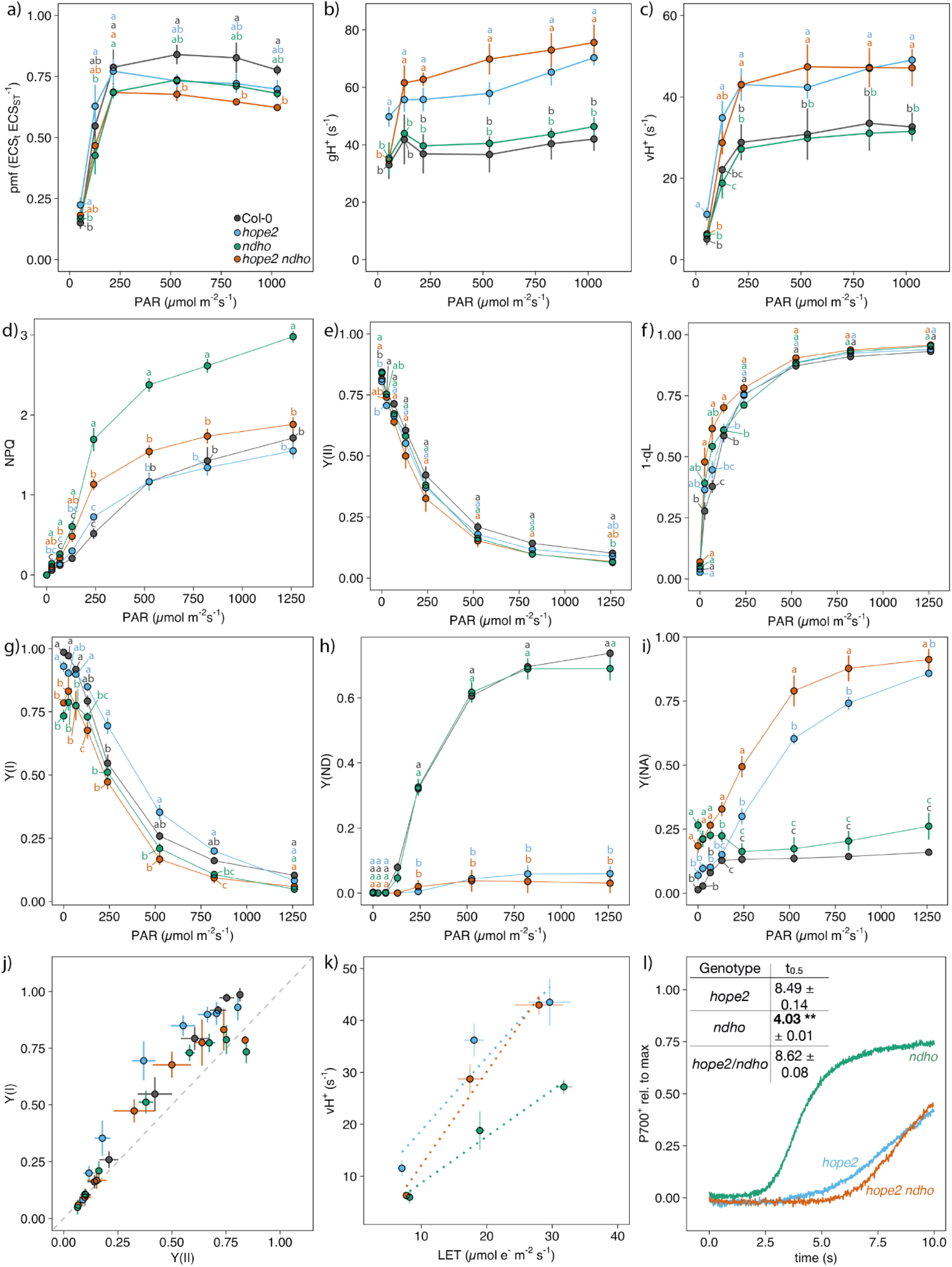
Photosynthetic parameters and CET measurements in *hope2 ndho*. a) proton motive force (pmf). b) proton conductance of ATP synthase (gH^+^). c) proton flux (vH^+^). d) nonphotochemical fluorescence quenching (NPQ). e) quantum yield of photosystem II Y(II). f) 1-qL (PSII acceptor-side limitation). g) quantum yield of photosystem I (Y(I)); h) donor-side limitation of PSI (Y(ND)). i) acceptor-side limitation of PSI (Y(NA)). j) Y(I) vs. Y(II). k) Proton flux vs. linear electron transfer. l) P700 oxidation during FR light, insert: t_0.5_ (s). Data points represent the average of 3-6 biological replicates ± SD. Colours represent genotypes analysed: Col-0 (black), *gl1* (grey), *hope2* (blue), *ndho* (green) and *hope2 ndho* (red). Different letters indicate statistical significance between genotypes at each light intensity, calculated from a Tukey HSD test, alpha = 0.05.

Since removal of NDH from the *hope2* background did not strongly affect the enhanced vH^+^ in *hope2* we next examined the effect of loss of PGR5. The Arabidopsis *pgr5* mutant has recently been shown to have point mutations in two separate genes encoding the PGR5 and PPT1 proteins, the latter of which seems to perpetuate long-term damage to PSI (Wada et al., 2021). Importantly for the conclusions of this work, however, the low pmf and Y(ND) phenotypes were shown to be associated with the PGR5 mutation alone, consistent with tDNA knock-out results in rice and a recent CRISPR-Cas9 *pgr5* mutant generated in *Arabidopsis* (Nishikawa et al., 2012; Penzler et al., 2022). Pmf and vH^+^ in the *hope2 pgr5* double mutant dropped significantly below *hope2*, except under very low light (Figure 4a,c), whereas gH^+^ was similar in *hope2* and *hope2 pgr5* (Figure 4b). In line with this, NPQ and Y(II) were lower in *hope2 pgr5* and 1-qL was higher compared to *hope2* (Figure 4d-f). Y(I) was also substantially lower at all light levels in *hope2 pgr5* compared to *hope2* (Figure 4g). Y(ND) remained low in the *hope2 pgr5* double mutant, similar to the respective single mutants (Figure 4h) and Y(NA) in *hope2 pgr5* was similar to *pgr5* and elevated with respect to *hope2* (Figure 4i). CET measured via the relationship between vH^+^ and LET or Y(I) and Y(II) was significantly less steep in *hope2 pgr5* compared to *hope2* and resembled WT (Figure 4j, k). In line with this, FR-induced P700 oxidation in *hope2 pgr5* was also much faster than in *hope2*, indicating that the increased CET is abolished and consistent with this the t_0.5_ was significantly lower than *hope2* (p<0.0001) (Figure 4l). Overall, these data show that in *hope2*, PGR5-dependent CET is required to maintain WT-level pmf by increasing vH^*+*^, whereas NDH-dependent CET only makes a significant contribution to pmf in the *hope2* background at lower light intensities.

**Figure 4:**
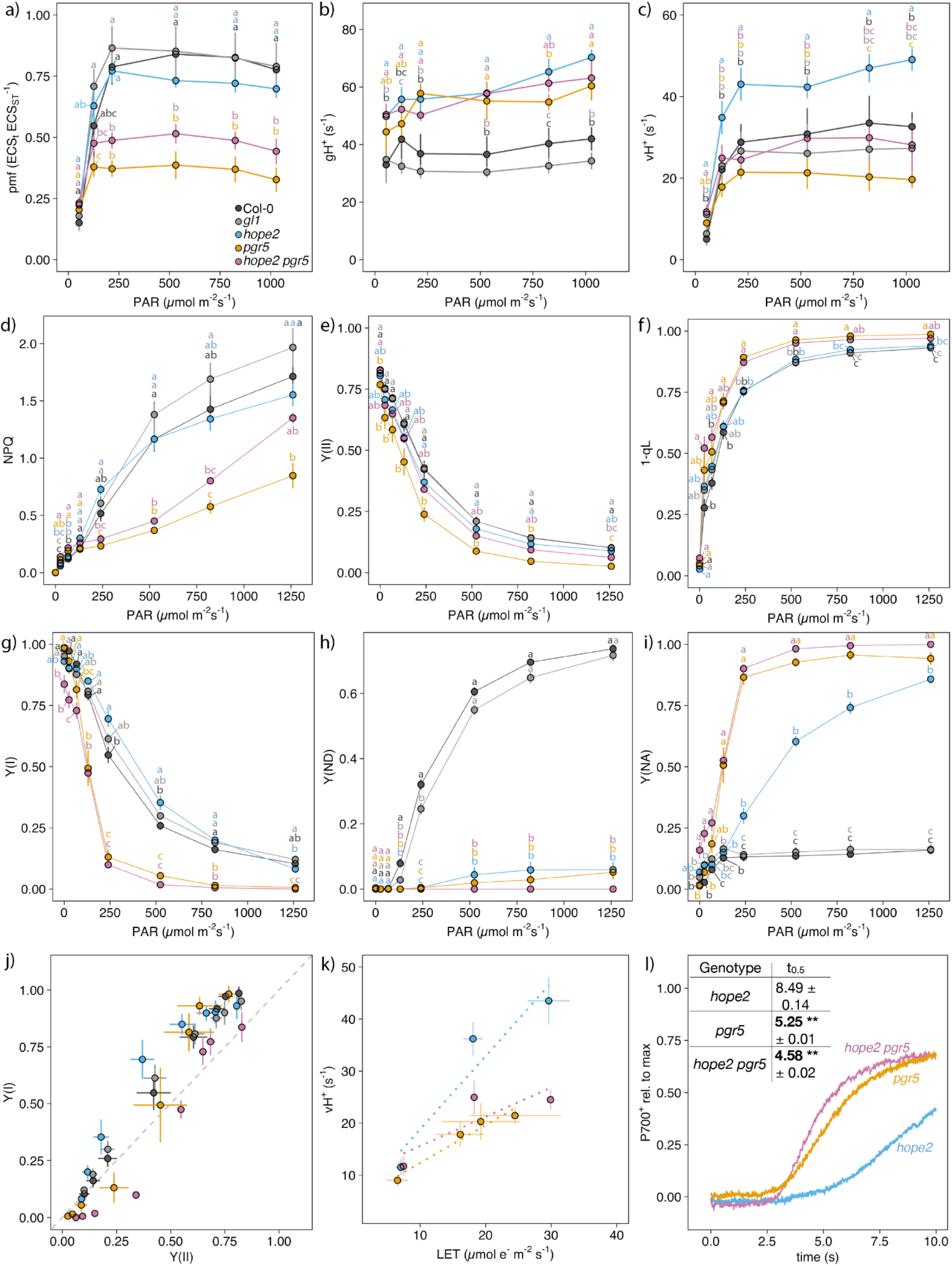
Photosynthetic parameters and CET measurements in *hope2 pgr5*. a) proton motive force (pmf). b) proton conductance of ATP synthase (gH^+^). c) proton flux (vH^+^). d) nonphotochemical fluorescence quenching (NPQ). e) quantum yield of photosystem II Y(II). f) 1-qL (PSII acceptor-side limitation). g) quantum yield of photosystem I (Y(I)); h) donor-side limitation of PSI (Y(ND)). i) acceptor-side limitation of PSI (Y(NA)). j) Y(I) vs. Y(II). k) Proton flux vs. linear electron transfer. l) P700 oxidation during FR light, insert: t_0.5_ (s). Data points represent the average of 3-6 biological replicates ± SD. Colours represent genotypes analysed: Col-0 (black), *gl1* (grey), *hope2* (blue), *ndho* (green) and *hope2 pgr5* (pink). Different letters indicate statistical significance between genotypes at each light intensity, calculated from a Tukey HSD test, alpha = 0.05.

### Photosynthetic control is absent in *hope2* despite maintenance of WT levels of ΔpH

The maintenance of WT levels of pmf in *hope2* due to increased PGR5-dependent CET raised the question of why NPQ is WT-like, while Y(ND) is *pgr5*-like? One possibility is that the pmf is differently partitioned between the ΔpH and ΔΨ components in *hope2*, which given the differing reported sensitivity of NPQ and Y(ND) to ΔpH might explain their contrasting responses (Horton et al., 1991; Nishio and Whitmarsh, 1993). Indeed, our proteomic data shows a increase in the relative abundance of the putative H^+^/K^+^ thylakoid antiporter KEA3, which could modify the ΔpH/ΔΨ partitioning of pmf in this mutant (Supplemental Figure S1). To test these ideas further we first confirmed that NPQ in *hope2* was of the ΔpH-dependent rapidly-relaxing qE type rather than photoinhibitory or sustained qI-type quenching (Supplemental Figure S4). Next we utilised the ECS partition method to assess the relative ΔpH and ΔΨ contributions to pmf. Previous ECS partitioning data suggested *hope2* may have a slightly lower ΔpH contribution to pmf (Takagi et al., 2017). However, this method has recently been called into question due to the overlapping absorption changes associated with qE which lead to overestimation of ΔΨ contributions to pmf, particularly when zeaxanthin synthesis is incomplete (Wilson et al., 2021). We thus compared using the partition method the pmf composition under conditions where zeaxanthin synthesis was incomplete (increasing light intensity every 20 seconds) versus complete (decreasing light intensity following 10 minutes illumination at 1421 µmol photons m^-2^ s^-1^). The results showed that pmf takes longer to establish in *hope2* consistent with the data in Figure 1c, as a result the apparent ΔΨ contribution to pmf is larger in *hope2* than in the WT (Figure 5a, b). In contrast, once pmf is established after 10 minutes of high light little difference in either the extent of pmf or the relative ΔΨ versus ΔpH contribution was observed between *hope2* and the WT (Figure 5b, c). In line with this, while NPQ is smaller in *hope2* during induction compared to the WT, once pmf is fully established in *hope2* NPQ reaches WT levels (Figure 5d). Yet, in spite of this Y(ND) remains much smaller and Y(NA) much larger in *hope2* (Figure 5e,f). Therefore, lower Y(ND) in *hope2* cannot be ascribed to lower ΔpH contribution to pmf.

**Figure 5:**
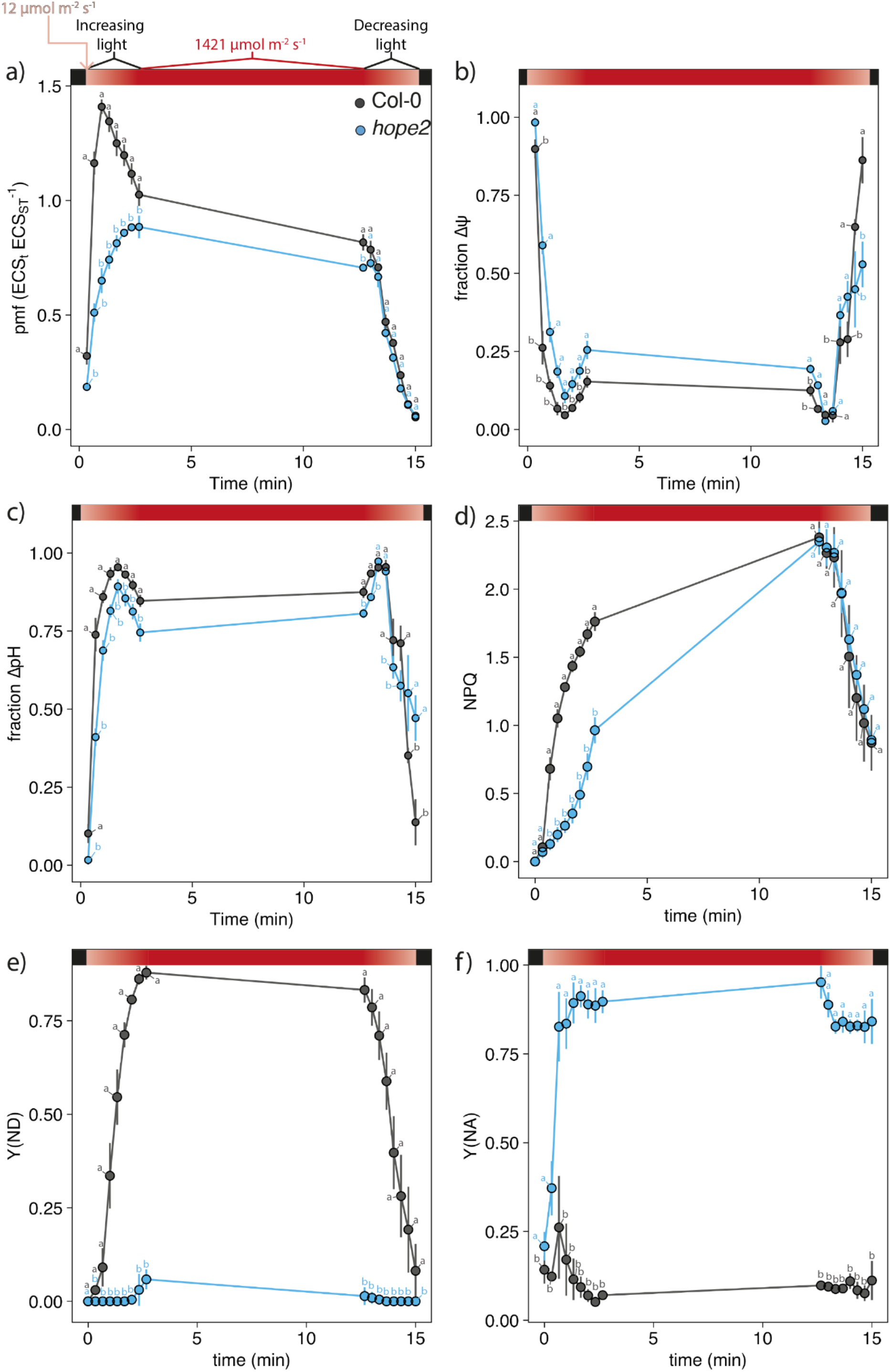
Response of Col-0 and *hope2* to rapidly increasing and decreasing light intensities. a) Proton motive force. b) Fraction of Δψ contribution. c) Fraction of ΔpH contribution. d) Nonphotochemical fluorescence quenching (NPQ). e) Donor-side limitation of PSI (Y(ND)). f) Acceptor-side limitation of PSI (Y(NA)). Rapid changes from low (12 µmol m-s s-1) to high (1421 µmol m-s s-1) light are indicated at the top of panel a by a red colour gradient. Light levels were increased every 20 sec, kept at high light for 10 min and decreased every 20 sec. Data points represent the average of 3 biological replicates ± SD. Colours represent genotypes analysed: Col-0 (black), *hope2* (blue). Different letters indicate statistical significance between genotypes at each light intensity, calculated from a Tukey HSD test, alpha = 0.05.

## Discussion

### *hope2* can maintain CO_2_ assimilation through increased CET

Proton motive force is harnessed for the production of ATP by ATP synthase, while its major ΔpH component also plays an important regulatory role triggering qE and Y(ND). *Hope2*, a recently described G134D γ1-subunit mutant in Arabidopsis was particularly interesting because it showed a similar phenotype compared to *pgr5* with respect to the loss of photosynthetic control and high gH^+^ but key differences with respect to qE and CO_2_ assimilation (Munekage et al., 2004; Takagi et al., 2017). Thus, ATP synthase regulation and CET may play distinct roles in the regulation of photoprotection and ATP/NADPH balance. Here we show that despite high gH^+^, *hope2* is able to maintain pmf at WT levels through increased vH^+^. The normal pmf levels we observe in *hope2* are a key point of difference with the previous study by Takagi *et al*., (2017). In this previous study pmf was more variable, being lower than WT under low O_2_ or ambient CO_2_ without pre-illumination and similar following pre-illumination. We found these differences could be explained by the slower establishment of pmf in *hope2* (Figs 1c, d, 5a). Comparison of the respective quantum yields of PSI and PSII showed increased electron transfer rate through PSI in *hope2* compared to WT consistent with enhanced CET (summarised in Figure 6). CET is notoriously difficult to measure because it produces no net product and utilises a common set of spectroscopically active-redox carriers with LET. Nonetheless, careful comparison on vH^+^ with the rate of LET demonstrates a steeper relationship in *hope2*, consistent with the phenotype seen in other high CET mutants previously described (Okegawa et al., 2005; Livingston et al., 2010a; Livingston et al., 2010b; Strand et al., 2015; Strand et al., 2017). A complementary method for assessing CET involves illumination monitoring the rate of P700 oxidation with FR. Consistent with the vH^+^/LET method the FR oxidation of PSI is slower in *hope2* confirming an increased rate of CET. The slower establishment of pmf in *hope2* (Figure 1c,d) seems to reflect the varying timescales for relaxation of ATP synthase and activation of CET. Thus, in the first few moments following illumination increases in pmf depend largely on the restricted gH^+^ in the WT. As the ATP synthase regulatory γ1-subunit thiol becomes gradually reduced in the light gH^+^ increases (Konno et al., 2012). Following this increase in gH^+^, vH^+^ must be increased if pmf is to be maintained. The maintenance of WT-levels of pmf likely explains the similar CO_2_ assimilation rates in *hope2* compared to the WT and is further evidence for an important role of CET in ensuring the optimal ATP/NADPH ratio.

**Figure 6:**
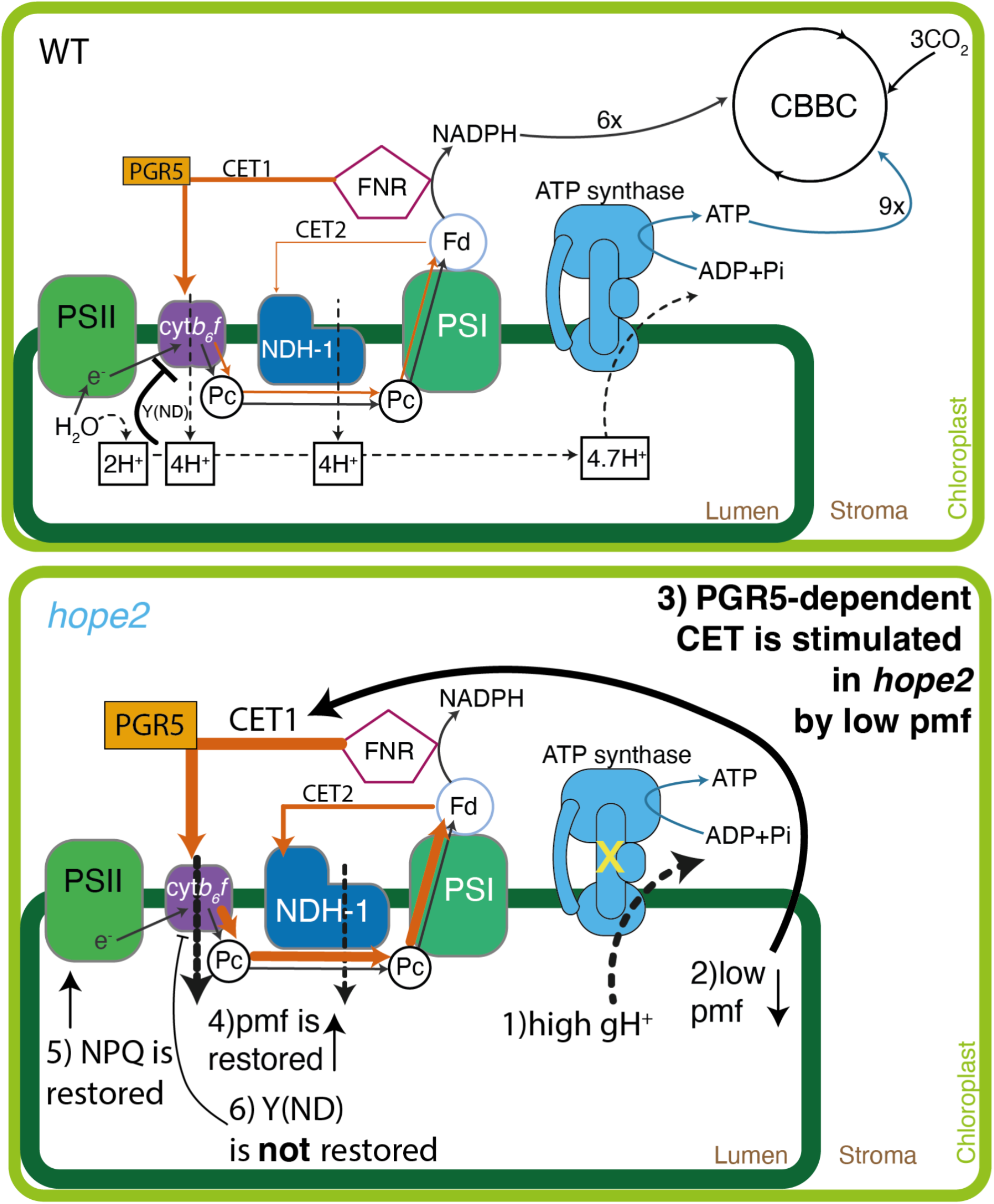
Proposed model of PGR5-dependent supercharged CET in *hope2*. In WT, CET is dominated by PGR5 to produce extra pmf and ATP. CET2 via NDH is only important at low light. Metabolic control of ATP synthase in *hope2* is disturbed, resulting in high gH^+^ and low pmf. In response, PGR5 levels are increased, and to a lesser extent, NDH levels, too. This increases CET1 and restores pmf and NPQ. However, Y(ND) is not restored, suggesting that metabolic control of ATP synthase is crucial for PSI photoprotection.

### Enhanced CET in *hope2* depends on the PGR5-dependent rather than NDH-dependent pathway

A proteomic comparison of the thylakoid membranes from *hope2* and WT plants showed that both NDH and PGR5 abundance was upregulated. We therefore constructed the double mutants *hope2 pgr5* and *hope2 ndho*, to understand the respective contributions of the two pathways to increased CET in *hope2*. The steeper vH^+^ versus LET and Y(I) versus (YII) slopes and slower FR-driven P700 oxidation seen in *hope2* were lost in *hope2 pgr5* double mutant, and it suffered from the more extreme PSI acceptor side limitation (Y(NA)) seen in *pgr5*. In contrast the phenotype of the *hope2 ndho* double mutant was less dramatic, with a similar rate of P700 oxidation to *hope2*, with only a slight decrease in the vH^+^/LET slope seen at low light and although there was an increase in Y(NA) compared to *hope2* this was less severe than in *hope2 pgr5*. The predominant dependence of *hope2* on the PGR-dependent CET1 pathway is key point of difference compared to other previously characterised high cyclic electron flow (*hcef*) mutants, which depended on the NDH-dependent CET2 pathway (Livingston et al., 2010a; Livingston et al., 2010b). The factors regulating CET1 and CET2 have yet to be fully elucidated (Yamori and Shikanai, 2015; Yamori et al., 2016), though H_2_O_2_ signalling and NADPH/ NADP^+^ redox balance have been recently implicated in control of the NDH and PGR5 pathways respectively (Strand et al., 2015; Strand et al., 2016). However, since Y(NA) is high in *hope2*, both signals would be expected, consistent with the increased abundance of both PGR5 (0.56-fold increase) and NDH subunits (0.21-0.41-fold increase) we find in this mutant. In contrast, quantification of NDH levels by immunoblotting in *hcef1* revealed a 15-fold increase in NDH and a 50% decrease in PGR5 (Livingston et al., 2010a). Comparison of *hope2* and *hcef1* reveals a large difference in gH^+^ between the mutants. Therefore, if pmf can be restored by combination of ATP synthase gH^+^ downregulation and increases in NDH-dependent CET2, it may negate the need for upregulation of PGR5. It may be significant in this regard that we see the greatest contribution of the NDH-pathway (H^+^/e^-^ ratio of 4) in *hope2* under low light, where it is less thermodynamically limited by backpressure from the pmf (Strand et al., 2017). This is in line with previous work showing *ndho* and *crr* mutants show more significant phenotypes in low light situations (Yamori et al., 2011; Wang et al., 2015; Yamori et al., 2015; Yamori and Shikanai, 2015). In contrast, the lower efficiency PGR5 pathway (H^+^/e^-^ ratio of 2) may be preferred in high light, consistent with the stronger phenotype of *pgr5* plants under such conditions (Munekage et al., 2004). Therefore, a combination of simple competition for excess electrons at the PSI acceptor side and thermodynamic constraints on turnover may determine which CET pathway is favoured in particular circumstances. Irrespective, the isolation of a high CET mutant that depends on PGR5 is significant since the steady-state contribution of CET1 is generally estimated to be low (<13% of LET) and difficult to measure. *Hope2* is therefore a useful tool for future research on PGR5-dependent CET1.

### High CET in the absence of ATP synthase regulation fails to restore photosynthetic control

Previously, repetitive flash treatment showed that PSI in *hope2* was, similar to *pgr5*, susceptible to photoinhibition (Takagi et al., 2017). However, it was unclear whether the *pgr5* phenotype was primarily due to loss of gH^+^ control or loss of CET1(Avenson et al., 2005; Yamamoto and Shikanai, 2020). While the maintenance of WT-levels of pmf through enhanced CET in *hope2* allowed normal qE-levels to develop, Y(ND) remained virtually absent (summarised in Figure 6). *Prima facie* this suggests that gH^+^ regulation of ATP synthase is crucial for photosynthetic control. Accordingly, even strongly enhanced PGR5-dependent CET1 does not protect against PSI photoinhibition in *hope2*. The failure of CET to protect PSI is consistent with recent results showing that both CET1 and CET2 do not act as photoprotective electron sinks in the absence of other mechanisms of acceptor side regulation of PSI (Rantala et al., 2020). Since photosynthetic control relies on the low lumenal pH-induced slowdown of PQH_2_ oxidation by the Rieske iron-sulphur protein of cyt*b*_6_*f (Nishio and Whitmarsh, 1993; Jahns et al., 2002)*, the most logical explanation for loss of Y(ND) is a loss of ΔpH in *hope2*. Previously, a difference in relative partitioning of pmf into ΔΨ and ΔpH compared to the WT was found in *hope2* (Takagi et al., 2017). However, we traced these apparent differences in partitioning to a slower establishment of pmf in *hope2*, which leads to increased overlap with the qE related absorption changes as described for the *npq1* mutant lacking zeaxanthin (Wilson et al., 2021). Once pmf is established in *hope2*, and presumably zeaxanthin synthesis is completed, then no major differences in the amplitude of ΔpH between *hope2* and the WT are present. Therefore, changes in ΔpH are not the cause of the low photosynthetic control phenotype in *hope2*. These data mirror similar reports in *pgr5* plants overexpressing the *Chlamydomonas rheinhardtii* plastid terminal oxidase 2 (PTOX2) protein and the FNR antisense mutant of tobacco, both of which showed normal qE but lacked photosynthetic control ((Hald et al., 2008; Zhou et al., 2021). One possibility is that just the relationship between qE and ΔpH is modified by the xanthophyll cycle de-epoxidation state (Rees et al., 1989; Horton et al., 1991), so the relationship between Y(ND) and ΔpH is regulated by the metabolic state of the stroma. Interestingly, both Y(ND) and gH^+^ are restored in *pgr5* in the presence of the artificial PSI electron acceptor methyl viologen (Munekage et al., 2002; Wang et al., 2018) or via transgenic expression of *Physcomitrella patens* Flv proteins (Yamamoto et al., 2016). Therefore, it may be that an oxidised PSI acceptor pool is required to trigger both regulatory processes.

### Conclusion

Our data have clarified the respective importance of proton influx and efflux control in photosynthetic regulation. We found ATP synthase gH^+^ regulation is indispensable for photosynthetic control even when CET can maintain pmf to ensure an optimal ATP/NADPH ratio and qE. This work highlights the interconnectedness and mutual dependence of the various photoprotective regulatory mechanisms in addition to the remarkable ability of the photosynthetic apparatus to preserve pmf via molecular plasticity in thylakoid protein composition.

## Material and Methods

### Plant material and growth condition

Arabidopsis mutants *hope2* and the WT background Col-0 and *pgr5* and the WT background *gl1* were grown in a controlled-environment chamber for at least 6 weeks at 21/15 °C day/night, 60% rel. humidity with an 8-hour photoperiod at a light intensity of 200 µmol photons m^-2^ s^-1^. Double mutants were generated by crossing *hope2* with either *pgr5* or *ndho*. Seeds from successful crosses were sown and allowed to self-fertilise, before genotyping via sequencing and western blotting to verify homozygosity.

### Chlorophyll fluorescence and *in situ* P700 absorption spectroscopy

A Dual-KLAS-NIR photosynthesis analyser (Heinz Walz GmbH, Effeltrich, Germany) was used for pulse-amplitude modulation chlorophyll fluorescence measurements and P700 absorption spectroscopy in the near-infrared (Klughammer and Schreiber, 2016; Schreiber and Klughammer, 2016). After plants had dark-adapted for at least 1 h, four pairs of pulse-modulated NIR measuring beans were zeroed and calibrated before each measurement. For each genotype, one leaf was used to generate differential model plots according to manufacturer’s protocol, which were used for online deconvolution to determine redox changes of P700. Prior to each measurement, maximum oxidation of P700 was determined by using the pre-programmed NIRmax routine. This consisted of a 3 s pulse of actinic light on top of which a 30 ms multiple turnover flash (MT) was given after 800 ms, followed by 4 s of darkness and 10 s of 255 µmol photons m^-2^ s^-1^ FR light and a MT at the end to achieve full oxidation of P700. NIRmax values were determined by using the pre-programmed “Get Max-Values” option. Dark-fluorescence (Fo) and maximal fluorescence (Fm) were determined prior to light or induction curves. Photosynthetic parameters were determined by using measuring beam intensities of 20 µmol photons m^-2^ s^-1^ and 14 µmol photons m^-2^ s^-1^ for chlorophyll fluorescence and P700 redox changes, respectively and a 18,000 µmol photons m^-2^ s^-1^ saturating pulse. Photosynthetic parameters were calculated as follows: Y(II) = (Fm’-F)/Fm’, NPQ = (Fm-Fm’)/Fm’, Y(I) = (Pm’-P)/Pm, Y(NA) = (Pm-Pm’)/Pm, Y(ND) = (P-P_0_)/Pm. For light curves, measurements were taken after 5 min at each light intensity. For induction curves, AL intensity was set to 169 µmol photons m^-2^ s^-1^ and measurements were taken at multiple time points after AL was turned on. To determine P700 oxidation for CET determination, leaves were exposed to a weak measuring light for 30 s, followed by a MT. FR light was then turned on for 20 s. Half-time of P700 oxidation was determined by fitting an allosteric sigmoidal function (Graphpad Prism, 9.1.1).

### Electrochromic shift measurements

Electrochromic shift was measured using a Dual-PAM analyser with a P515/535 emitter /detector module (Heinz Walz GmbH, Effeltrich, Germany) (Klughammer et al., 2013). Plants were dark-adapted for at least 1 h prior to measurements. Proton motive force (pmf) was calculated from the decay of the P515 signal when AL was turned off, by fitting a single exponential decay to the first 300 ms in the dark to determine the span of the signal decay (ECS_t_). pmf was normalised by dividing ECS_t_ by the magnitude of a 50 µs ST flash applied prior to account for leaf thickness and chloroplast density (Takizawa et al., 2007; Livingston et al., 2010a; Wang et al., 2015; Takagi et al., 2017). The proton conductance gH^+^ was calculated as the inverse of the decay time constant *τ*_ECS_ of the single exponential decay and proton flux was calculated as vH^+^= pmf x gH^+^ (Baker et al., 2007).

### Leaf infiltration

Dark-adapted leaves were vacuum-infiltrated with 30 µM DCMU and 100 µM methyl viologen buffered in 20 mM Hepes - pH 7.5, 150 mM sorbitol and 50 mM NaCl. Leaves were dark-adapted for 10 min between infiltration and measurements.

### Gas exchange

CO_2_-response (ACi) and light-response curves (AQ) were measured using the infrared gas analyzer system 6400-XT (LiCOR Biosciences, Lincoln, NE, USA). Prior to measurements, plants were exposed to 400 ppm reference CO_2_, 1500 µmol photons m^-2^ s^-2^ light at 25 °C and ca. 50% relative humidity for at least 10 min until steady state was reached and stomata were wide open with a Ci/Ca of >0.7. For ACi curves, data was logged at various CO_2_ concentrations after 90-120 s using the following sequence of reference CO_2_ concentrations, as recommended by (Busch, 2018): 400, 350, 300, 250, 200, 150, 100, 50, 400, 400, 450, 500, 650, 800, 1000, 1250. For AQ curves, sample CO_2_ was set to 390 ppm and data was logged after a minimum of 3 min at each light intensity, using the following sequence of light intensities: 1500, 1000, 750, 500, 300, 200, 150, 50, 25, 10, 0. Reference and sample analysers were matched prior to logging the data. Maximum Rubisco activity (Vc,max) and maximum electron transport rate used in RuBP regeneration (Jmax) were fitted using the FvCB model and the Plantecophys package in R (Duursma, 2015).

### Thylakoid isolation

Thylakoids were isolated from Arabidopsis plants 2-3 hours into the photoperiod. Plants were blended in ice-cold medium containing 50 mM sodium phosphate pH 7.4, 5 mM MgCl_2_, 300 mM sucrose and 10 mM NaF. The homogenate was then filtered twice through two layers of muslin cloth. The filtrate was then centrifuged for 15 min at 3750 rpm at 4 °C. The chloroplast pellets were then resuspended in 5 mM MgCl_2_, 10 mM Tricine pH 7.4 and 10 mM NaF. After 1 min on ice, a medium containing 5 mM MgCl2, 10 mM Tricine pH 7.4, 400 mM sucrose and 10 mM NaF was added. The broken chloroplasts were centrifuged for 15 min at 3750 rpm at 4 °C, and thylakoid pellets were resuspended in 10 mM sodium phosphate pH 7.4, 5 mM MgCl2, 5 mM NaCl and 100 mM sucrose. Resuspended thylakoids were centrifuged again, and pellets were resuspended in 1 mL of the same medium.

### Quantitative proteomic analysis of thylakoid membranes

Thylakoid membrane proteins were solubilized and digested with a combination of endoproteinase Lys-C and trypsin in 1% (w/v) sodium laurate, 100 mM triethylammonium bicarbonate pH 8.5 with additional sample processing and analysis by nano-flow liquid chromatography-mass spectrometry as previously described (Flannery et al., 2021). MaxQuant v. 1.6.3.4 (Cox and Mann, 2008) was used for mass spectral data processing and protein identification with the iBAQ (Schwanhäusser et al., 2011) label-free quantification option selected and other parameters as previously specified (Flannery et al., 2021). iBAQ abundance scores subjected to statistical analysis using a modified Welch’s t-test as implemented in Perseus v. 1.6.2.3 Protein identification and label-free quantification were performed using the MaxLFQ algorithm embedded within FragPipe (v. 16.0) (Yu et al., 2021). The ‘match-between-runs’ option was selected and all other parameters were as per default. Total intensities were normalised against the sum of all identified proteins. Not all proteins were identified by mass spectrometry, therefore only proteins where >75% of replicates where identified were selected. The normal distribution of data was verified in MaxQuant (Cox and Mann, 2008), imputed and log2 transformed. This data was then expressed as fold-change relative to WT, where p >0.05 was significantly different. The proteomics data have been deposited to the ProteomeXchange Consortium via the PRIDE partner repository (http://proteomecentral.proteomexchange.org) with the dataset identifier PXD033007.

### Statistical analysis

Statistical analysis was performed using Graphpad Prism, 9.1.1, using a two-sided t-test (alpha = 0.05) and the Tukey HSD test (alpha = 0.05) in R. The asterisks always indicate significant differences between the Col-0 & *hope2* and *gl1 &* pgr5. Different letters indicate significant differences.

## Acknowledgements

We would like to thank Prof. Toshiharu Shikanai for the gift of *pgr5* seeds, Prof. Chikahiro Miyake for the gift of *hope2* seeds and Prof. Eva-Mari Aro for *ndho* seeds. M.P.J. and S.A.C. acknowledge funding from the Leverhulme Trust grant RPG-2019-045 and M.P.J. from the Biotechnology and Biological Sciences Research Council (BBSRC) grant BB/V006630/1. For the purpose of open access, the author has applied a Creative Commons Attribution (CC BY) licence to any Author Accepted Manuscript version arising.

## Author contributions

G.E.D, M.S.P, P.J.J and N.Z performed research, S.A.C and M.P.J supervised research, G.E.D analysed the data and G.E.D and M.P.J wrote the paper. All authors approved of the manuscript prior to submission.

---

**Figure S1:**
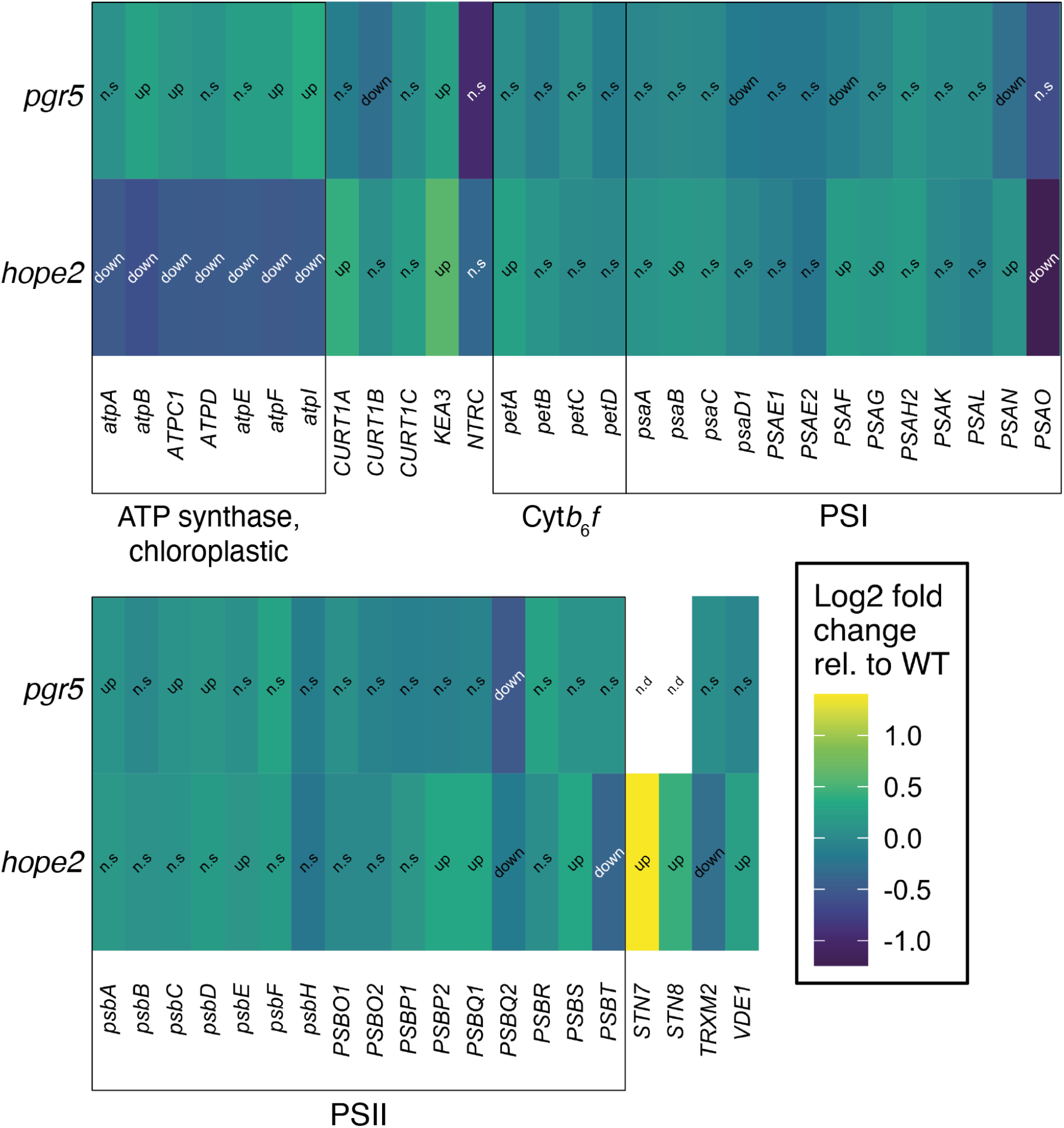
Abundance of photosynthetic proteins. Heat map log2 fold change relative to WT in hope2 and pgr5. “Up” indicates significant upregulation of protein, “down” indicates significant downregulation, “n.s.” indicates no significant change relative to WT and “n.d” indicates not detected.

**Figure S2:**
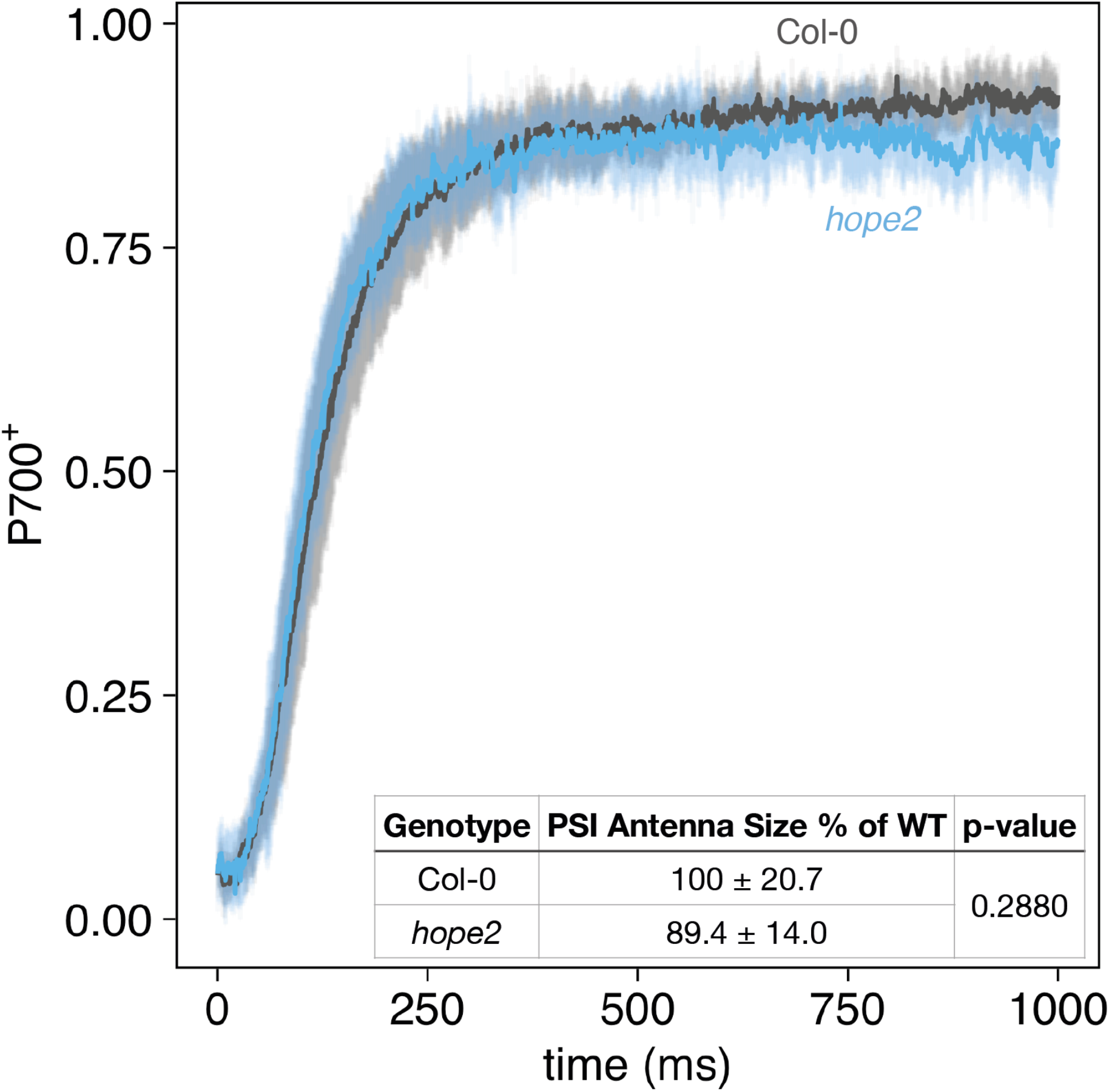
Determination of PSI antenna size in Col-0 and hope2. P700 oxidation was measured on intact leaves infiltrated with 30 µM DCMU and 100 µM methyl viologen to create a donor-side limitation and remove any acceptor-side limitation.

**Figure S3:**
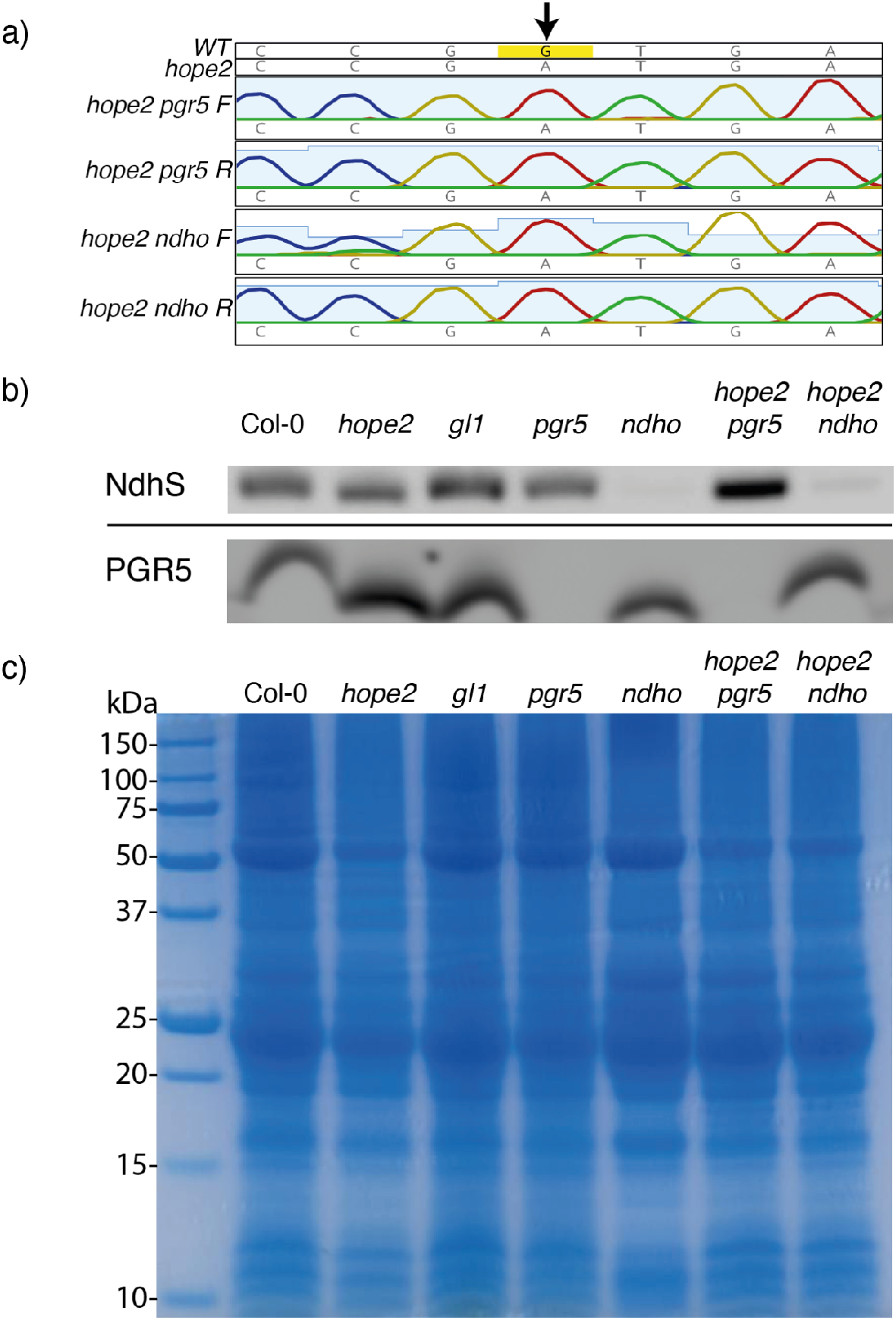
Verification of homozygosity of double crosses. a) Sequencing results of *hope2 pgr5* and *hope2 ndho* double crosses using forward (ACTTCCTCACCTCCTTCACG) and reverse (AATTTCCCTTCTTGCCCACG) primers to verify homozygosity of the A to G mutation (arrow). b) Western blot of isolated thylakoids using the anti-NdhS and anti-PGR5 antibodies to verify double crosses. c) Coomassie Brilliant Blue stain of SDS-PAGE gel used for Western blot in (b) to show equal amounts of protein were loaded.

**Figure S4:**
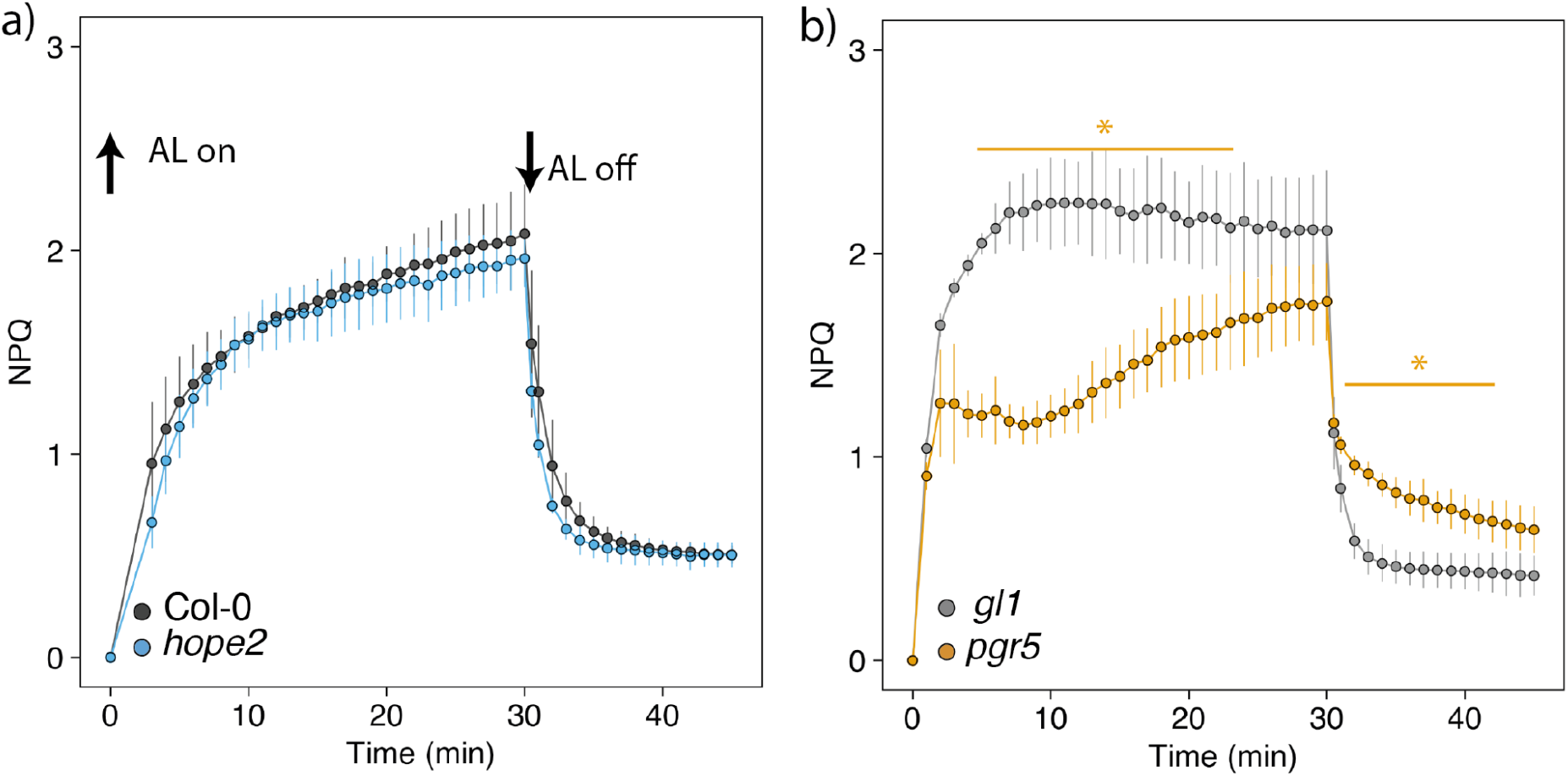
Determination of rapidly relaxing NPQ (qE). NPQ was recorded every minute during 30 min of 981 µmol m^-2^ s^-1^ AL and for 15 min after AL off. Data points represent the average of 3 biological replicates ± SD. Asterisks indicate a significant difference between (a) *hope2* (blue) and Col-0 (black) and (b) *pgr5* (orange) and *gl1* (grey), calculated from a two-sided t-test (alpha = 0.05, * p<0.05, ** p<0.01). Different letters indicate statistical significance between genotypes at each light intensity, calculated from a Tukey HSD test, alpha = 0.05.

